# Theta-burst stimulation over primary somatosensory cortex modulates tactile acuity of tongue

**DOI:** 10.1101/2024.06.17.599457

**Authors:** Ding-lan Tang, Mark Tommerdahl, Caroline A. Niziolek, Benjamin Parrell

## Abstract

**Background:** Emerging studies in humans have established the modulatory effects of repetitive transcranial magnetic stimulation (rTMS) over primary somatosensory cortex (S1) on somatosensory cortex activity and perception. However, to date, research in this area has primarily focused on the hand and fingers, leaving a gap in our understanding of the modulatory effects of rTMS on somatosensory perception of the orofacial system and speech articulators.

**Objective:** The present study aimed to examine the effects of different types of theta-burst stimulation—continuous TBS (cTBS), intermittent TBS (iTBS), or sham—over the tongue representation of left S1 on tactile acuity of the tongue.

**Methods:** In a repeated-measures design, fifteen volunteers participated in four separate sessions, where cTBS, iTBS, sham, or no stimulation was applied over the tongue representation of left S1. Effects of TBS were measured on both temporal and spatial perceptual acuity of tongue using a custom vibrotactile stimulator.

**Results:** CTBS significantly impaired spatial amplitude threshold at the time window of 16-30 minutes after stimulation, while iTBS improved it at the same time window. The effect of iTBS, however, was smaller than cTBS. In contrast, neither cTBS nor iTBS had any effect on the temporal discrimination threshold.

**Conclusions:** The current study establishes the validity of using TBS to modulate somatosensory perception of the orofacial system. Directly modifying somatosensation in the orofacial system has the potential to benefit clinical populations with abnormal tactile acuity, improve our understanding of the role of sensory systems in speech production, and enhance speech motor learning and rehabilitation.

**Highlights:**

- Theta-burst stimulation (TBS) can modulate somatosensation in the orofacial system
- cTBS over S1 impaired spatial acuity of tongue
- iTBS over S1 improved spatial acuity of tongue

## Introduction

Somatosensory input is essential for motor control. If somatosensory input is not properly processed both before and during movement, the resulting motor output exhibits abnormalities and/or inaccuracies (Riemann and Lephart, 2002; Cascio, 2010). Growing evidence indicates impairments in somatosensory function are a major contributor to the motor dysfunction commonly observed in neurologic injury or disorders such as Parkinson’s disease, cerebral palsy, dystonia, ataxia, etc. (Conte et al., 2013; Elbert et al., 1998; Konczak & Abbruzzese, 2013; Zarkou et al., 2020).

Primary somatosensory cortex (S1) plays a crucial role in somatosensory processing, and importantly, has demonstrated a high capacity for plastic change (Schaechter et al., 2006). Changes in response properties of S1 neurons and local excitatory/inhibitory circuitry in S1 have been observed in animals in response to manipulation of sensory experience (Blake et al., 2010; Cheetham et al., 2007) and repetitive intracortical microstimulation (Dinse et al., 1997; Heusler et al., 2000). In humans, emerging studies have used forms of repetitive transcranial magnetic stimulation (rTMS) as a tool to investigate somatosensory perception (see review, Tang et al., 2023). More recently, theta burst stimulation (TBS) has gained popularity and has been shown to modulate both somatosensory cortex activity and somatosensory perception (Kumar et al., 2019; Ragert et al., 2008; Rai et al., 2012). TBS can be applied intermittently (iTBS) or continuously (cTBS), resulting in either facilitatory or inhibitory effects, respectively (Huang et al., 2005). For example, an application of iTBS over S1 has been shown to significantly improve the tactile perception of the left index finger (e.g. Conte et al, 2012), whereas cTBS over S1 induced a pronounced impairment of tactile perception in the hand (e.g. Conte et al, 2012; Rai et al., 2012).

Previous studies in this area have predominantly centered on the hand and fingers, resulting in a dearth of understanding of the modulatory effects of rTMS on the orofacial system and speech articulators. Orofacial somatosensation plays an important role in speech motor learning (Darainy et al., 2019; Feng et al., 2011; Ito et al., 2016), and recent computational simulation work suggests somatosensory input is as important for accurate speech production as it is for limb and posture control (Parrell et al. 2019). TMS has potential to both enhance and impair all aspects of orofacial somatosensation, an improvement over existing methods like anesthesia, which can only block tactile sensation. Thus, modulating somatosensory activity and perception of the orofacial system and speech articulators with TBS has promise both in basic science work to improve our understanding of the role of sensory systems in speech production and, potentially, to modulate speech motor learning and rehabilitation, especially for people with speech motor disorders. The present study aimed to examine the effects of different types of theta-burst stimulation (cTBS, iTBS, sham) over the tongue representation of left S1 on the tactile acuity of the tongue.

## Method

### Subjects

Seventeen right-handed adult individuals were recruited. All participants reported no hearing problems and no personal or family history of seizures or other neurological disorders. Participants provided informed consent and were compensated monetarily for their participation. The Institutional Review Board of the University of Wisconsin-Madison approved the experimental protocol.

One participant withdrew after finishing the first visit for reasons unrelated to the study. One participant was excluded because we failed to localize their tongue representation in motor cortex (i.e. no clear tongue motor-evoked potentials (MEPs) could be observed under safe maximum TMS intensity). In total, data from 15 participants (8/7 females/males, 18-48 years, mean 25.3 ± 8.5) were further analyzed.

### Experimental procedure

The current study used a within-subject design. Each participant completed 4 sessions in which they underwent the same behavioral procedure (see Figure 1). There was at least one week between each session. During the first three visits, participants received cTBS, iTBS or sham stimulation over the left S1. The order was counterbalanced across participants. The fourth (control) visit involved only behavioral tasks without any brain stimulation.

**Figure 1.**
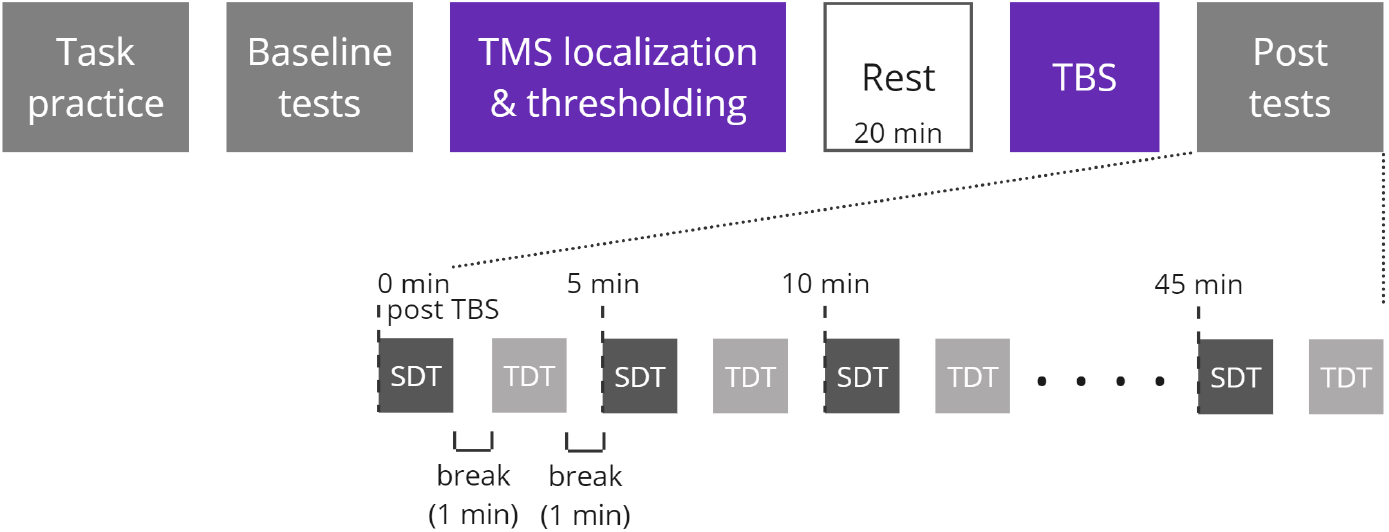
Experimental procedure of each visit. During each visit, participants received a different type of TBS (cTBS, iTBS, or sham) or no stimulation (control). Spatial amplitude discrimination threshold (SDT, dark grey) and temporal discrimination threshold (TDT, light grey) were estimated before stimulation (baseline tests) and every 5 minutes for 45 minutes following stimulation (post tests).

The timeline of each visit is depicted in Figure 1. Spatial amplitude discrimination threshold (SDT, ∼80s/block) and Temporal discrimination threshold (TDT, ∼100s/block) tasks were used to measure spatial and temporal tactile acuity of the tongue, respectively (see Task and Stimuli below for more detailed task information). Participants first completed a brief training to familiarize them with each task. Once performance criteria of the training trials were met (five consecutive correct responses), baseline (pre-TBS) tests began. Baseline tests consisted of 3 blocks of each task, with the tasks alternating. A 1-min break was given after each block. When the baseline tests were completed, participants received single-pulse TMS to localize S1 and determine the active motor threshold . TBS was delivered following a 20 minute break to allow the somatosensory cortex to recover from any potential disruption related to single-pulse TMS. Finally, post-TBS tests were performed every 5 minutes for 45 minutes following stimulation: participants completed 10 blocks of each task (SDT and TDT), alternating between the tasks with a 1-minute break in between each task.

### Task and Stimuli

A custom vibrotactile stimulator (Cortical Metrics) was used to measure both the temporal discrimination threshold and spatial amplitude discrimination threshold of the tongue. A similar device has previously been used to measure the temporal and spatial amplitude acuity of the hand (Lee et al., 2013; Rai et al., 2012; Tannan et al., 2005). The device consists of two plastic circular (3-mm diameter) probes located on the surface of a rotatory cylindrical disk. The distance between the two probes was set at approximately 2.4 cm to ensure participants could feel both probes, one on each side of the tongue.

During both tasks, participants were seated comfortably in a chair in front of the vibrotactile stimulator, which was placed on a height-adjustable stand set to a comfortable position. Participants positioned their tongue on the two probes and were instructed to try to maintain even pressure against both probes. During the 1-min break between tasks, participants were able to relax their tongue. They were instructed to reposition their tongue on the probes to get ready for the task at least 10 s before the next block (a countdown timer was shown on a computer monitor). Earplugs were provided to minimize the auditory cues resulting from the vibrating probes.

#### Spatial amplitude discrimination threshold (SDT) task

Each block of the SDT task consisted of 15 trials, with each block yielding a unique measurement of the participants’ discrimination threshold. For each trial, the two probes were vibrated simultaneously at 25 Hz for a duration of 500 ms. Participants were instructed to report which probe vibrated more intensely by clicking a computer mouse (left click = left probe is more intense, right click = right probe is more intense). The amplitude difference of vibration between the two probes was set at 100 μm for the first trial of each block and was subsequently varied with a step size of 20 μm. A 1-down 1-up rule was followed for the first 5 trials, and then adjusted to a 2-down 1-up rule for the rest of the trials (i.e., two correct responses decreased the amplitude difference by 20 μm while one incorrect response increased the amplitude difference by 20 μm). The amplitude of one probe (standard probe) remained constant at 200 μm throughout all trials. The location of the standard and test was randomized on a trial-by-trial basis. The inter-trial interval (ITI) was set at 4 s. The perceptual SDT for each block of the task was calculated as the average of the amplitude difference in the last five trials (Folger et al., 2008; Rai et al., 2012; Zhang et al., 2008).

#### Temporal discrimination threshold (TDT) task

Each block of the TDT task consisted of 21 trials, with each block yielding a unique measurement of the participants’ discrimination threshold. For each trial, the two probes were vibrated at 25 Hz for 40 ms at an amplitude of 300 μm. The interstimulus interval (ISI) between the first and second vibrating probe was set at 150 ms for the first trial of each block and was subsequently varied following a 1-down 1-up rule, with a step size of 15%. Participants were instructed to identify which probe vibrated first and respond as quickly as possible by making a mouse click (left click = left probe vibrated first, right click = right probe vibrated first) with their right hand. The first vibrating probe (left or right) was randomized on a trial-by-trial basis. The ITI was set at 3 seconds. The perceptual TDT for each block of the task was calculated as the average ISI measured from the last five trials (Lee et al., 2013; Nelson et al., 2012; Rai et al., 2012; Tommerdahl et al., 2008).

### Electromyography (EMG) recording

EMG was recorded using a BrainSight built-in 2-channel EMG device. To record EMG activity from the tongue muscles, two disposable EEG cup electrodes were mounted to a swimming nose clip, which was placed on the right side of the tongue blade (electrodes placed on dorsal and ventral surfaces) (Panouillères & Möttönen, 2018; Tang et al., 2021). The ground electrode was placed on the forehead for tongue EMG recordings.

### TMS and neuronavigation

TMS was delivered using Magstim 200 (single pulse) and Magstim Super Rapid Plus (TBS) stimulators with a D702 70-mm figure-of-eight coil (Magstim Company LTD, United Kingdom). A BrainSight neuronavigation system (Rogue Research, Montreal, Canada) was used to digitally register an MNI template brain (BrainSight default) with the TMS coil.

After baseline testing, single-pulse TMS (monophasic, Magstim 200) was used to localize the tongue representation in the left primary motor cortex (M1 hotspot), following a similar procedure as previously described (Tang et al., 2021). The coil was placed tangential to the skull, aiming to induce a current flow from posterior to anterior under the junction of the two wings of the figure-of-eight coil. The coil’s position was adjusted until robust tongue MEPs were consistently observed. After localization, biphasic single-pulse TMS (Magstim Super Rapid) was applied over the M1 hotspot to determine the active motor threshold, the minimum intensity at which TMS elicited MEPs with an amplitude of at least 200 μV on at least 5 out of 10 pulses when the tongue muscle was contracted at 20-30% of the maximum. Participants were trained to maintain the target level of muscle contraction using EMG feedback from the electrodes placed on the tongue, displayed visually on a computer monitor. S1 was initially localized by moving 2 cm posterior from M1 hotspot on the cortex as measured using Brainsight Neuronavigation (Tang et al., 2023). If single-pulse TMS over this region elicited MEPs from the tongue muscles, the coil was moved posteriorly until tongue MEPs were no longer elicited. The M1 hotspot, S1 location and active motor threshold were determined for each of the three TMS visits following the same procedure.

Following a 20-minute break after localization, participants received cTBS (bursts of 3 pulses at 50 Hz at a rate of 5 Hz for 40 seconds, 600 pulses in total), iTBS (bursts of 3 pulses at 50 Hz for 2 seconds, repeated every 10 s for a total of 190 seconds, 600 pulses) or sham stimulation over the tongue representation of left S1 at 80% of active motor threshold. The coil’s positioning and orientation (with the handle oriented 45° from the midline) was visually monitored throughout the whole stimulation. For sham stimulation, the coil was placed over S1 and rotated 90°, with the coil handle pointing vertically upward while still maintaining contact with the scalp.

## Data analysis

For both TDT and SDT, we defined the baseline threshold for each participant as the average of the discrimination thresholds estimated from the last two blocks of the baseline tests. We omitted the first block from the average because some participants showed large changes in thresholds from the first to the second block, likely due to increased familiarization. Threshold estimates were stable by the second baseline block, indicated by a lack of difference for either task between the second and third baseline blocks (paired t-test, *p* > .05 in all sessions). Baseline thresholds for each task across sessions (cTBS, iTBS, sham and control) were compared using repeated-measures ANOVAs. Post-TBS values were normalized by subtracting baseline thresholds for each task on a session-specific basis. During post-TBS tests, normalized discrimination thresholds were binned into three time windows (0-15 min, 16-30 min, and 30+ min after stimulation). We first assessed the significance of threshold changes in each task using two-tailed one-sample t-tests with a significance level of a = .05, comparing baseline-normalized thresholds in each of the three time windows against 0 (no change from baseline) for each session. To compare changes in thresholds across sessions, a single repeated-measures ANOVA was conducted for each task (SDT, TDT) with sessions (cTBS, iTBS, sham and control) and time windows (0-15, 16-30, 30+ min) as independent variables. In the event of an interaction, follow-up post hoc comparisons at each time window were conducted using a repeated-measures ANOVA across sessions. Further post-hoc pairwise comparisons between sessions at a particular time window (one-tailed t-tests with Bonferroni correction, given our prior hypothesis regarding the effects of different stimulations) were conducted only in the event of a significant main effect of this follow-up ANOVA.

All statistical analyses were performed in R (R Core Team). Repeated-measures ANOVAs and t-tests were performed with the rstatix package (Kassambara, 2021). Partial eta squared (η^2^) and Cohen’s d were calculated for repeated-measures ANOVAs and t-tests, respectively, to determine effect size for statistically significant effects.

## Results

The mean active motor thresholds (% stimulator output) for each stimulation were 38.3 ± 5.9 (cTBS), 38.7 ± 7.4 (iTBS) and 37.7 ± 5.9 (sham). There was no significant difference in active motor thresholds across the three stimulation sessions (F_(2, 28)_ = 1.25, p = .302).

### Spatial amplitude discrimination threshold (SDT) task

The mean spatial amplitude discrimination thresholds during the baseline phase were 34.1 ± 20.9 μm (cTBS), 43.8 ± 16.9 μm (iTBS), 37.9 ± 16.6 μm (sham), and 40.9 ± 21.8 μm (no stimulation control), with no significant difference in SDT across sessions (F_(3, 42)_ = 1.31, p = .283, Figure S1). To further evaluate the reliability of the SDT task, the intraclass correlation coefficient (ICC, using average measures) was calculated based on a 2-way mixed-effects model. The across-session ICC for baseline SDT was estimated to be 0.78, suggesting a good reliability of the spatial amplitude discrimination task (Koo & Li, 2016).

Stimulation with cTBS significantly impaired spatial amplitude discrimination threshold at the time window of 16-30 min after stimulation (+20.08 µm, t_14_ = 2.73, p = .016, d = 0.7), while iTBS improved it at the same time window (-8.07 µm, t_14_ = 2.16, p = .049, d = 0.56) (Figure 2A). The suppression and enhancement effects induced by cTBS and iTBS persisted numerically past 30 minutes after stimulation, though these effects did not reach statistical significance at this later time window (cTBS: +18.1 µm, t_14_ = 1.87, p = .082, d = 0.48; iTBS: -6.63 µm, t_14_ = 1.8, p = .093, d = 0.47). No significant change was observed at any post-TBS time window during the sham or control (no stimulation) sessions (p > .13 in all cases). Comparing between sessions (cTBS, iTBS, sham and control) revealed a significant interaction between session and time (F_(6,84)_ = 2.19, p = .05, partial η^2^ = 0.14). Follow-up ANOVAs at each time window showed that there were significant differences between sessions only at the time window of 16-30 min after stimulation (F_(3,42)_ = 5.49, p = .003, partial η^2^ = 0.28; other p > 0.05). Post-hoc tests revealed that the baseline-normalized thresholds after cTBS were significantly higher than those after iTBS (t_14_ = 4.22, p = .003, d = 1.09) and control (t_14_ = 2.72, p = .049, d = 0.7), but there was no significant difference between cTBS and sham (t_14_ = 1.68, p = .288). The baseline-normalized thresholds after iTBS, however, showed only marginally significant differences (after Bonferroni correction) when compared to the sham session (t_14_ = 2.04, p = .151, d = 0.53) and showed no significant difference when compared to the control session (t_14_ = 0.82, p = .848).

**Figure 2.**
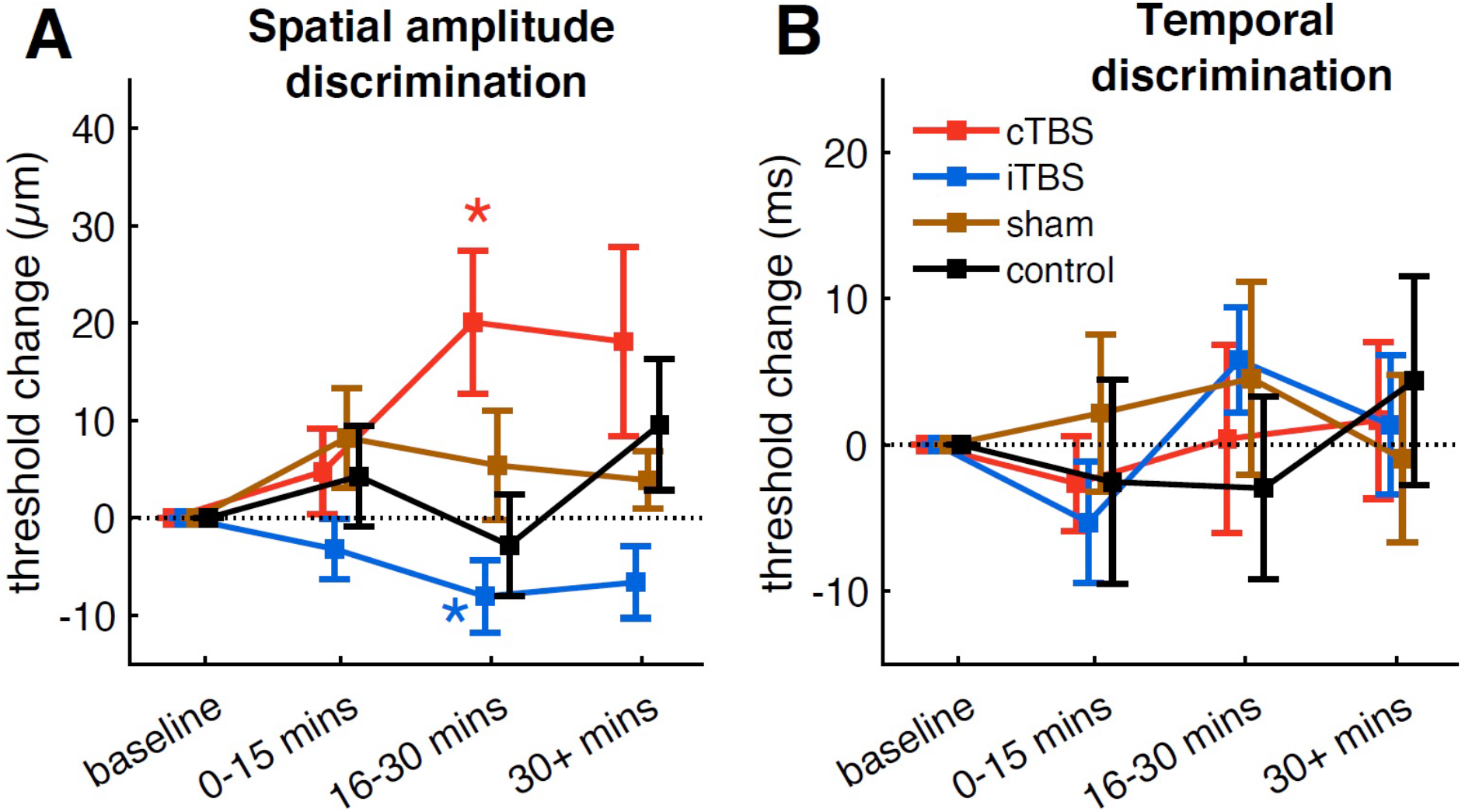
(A) Spatial amplitude discrimination thresholds (SDT); (B) Temporal discrimination thresholds (TDT). Baseline-normalized thresholds in the baseline, 0-15 min, 16-30 min, and 30 min after stimulation are shown for different groups. Error bars show standard error, and * indicates a significant change (p < .05) from baseline.

Group-averaged trial-by-trial spatial amplitude discrimination performance for each session is shown in Figure 3A. Performance across all time windows (baseline, 0-15 min, 16-30 min, and 30 min after stimulation) for all four sessions reached a plateau around trial 9 of 15, which is similar to previous experiments that measured spatial amplitude acuity of the hand (Rai et al., 2012). The effects of cTBS and iTBS on SDT occurred when performance had already reached the plateau.

**Figure 3.**
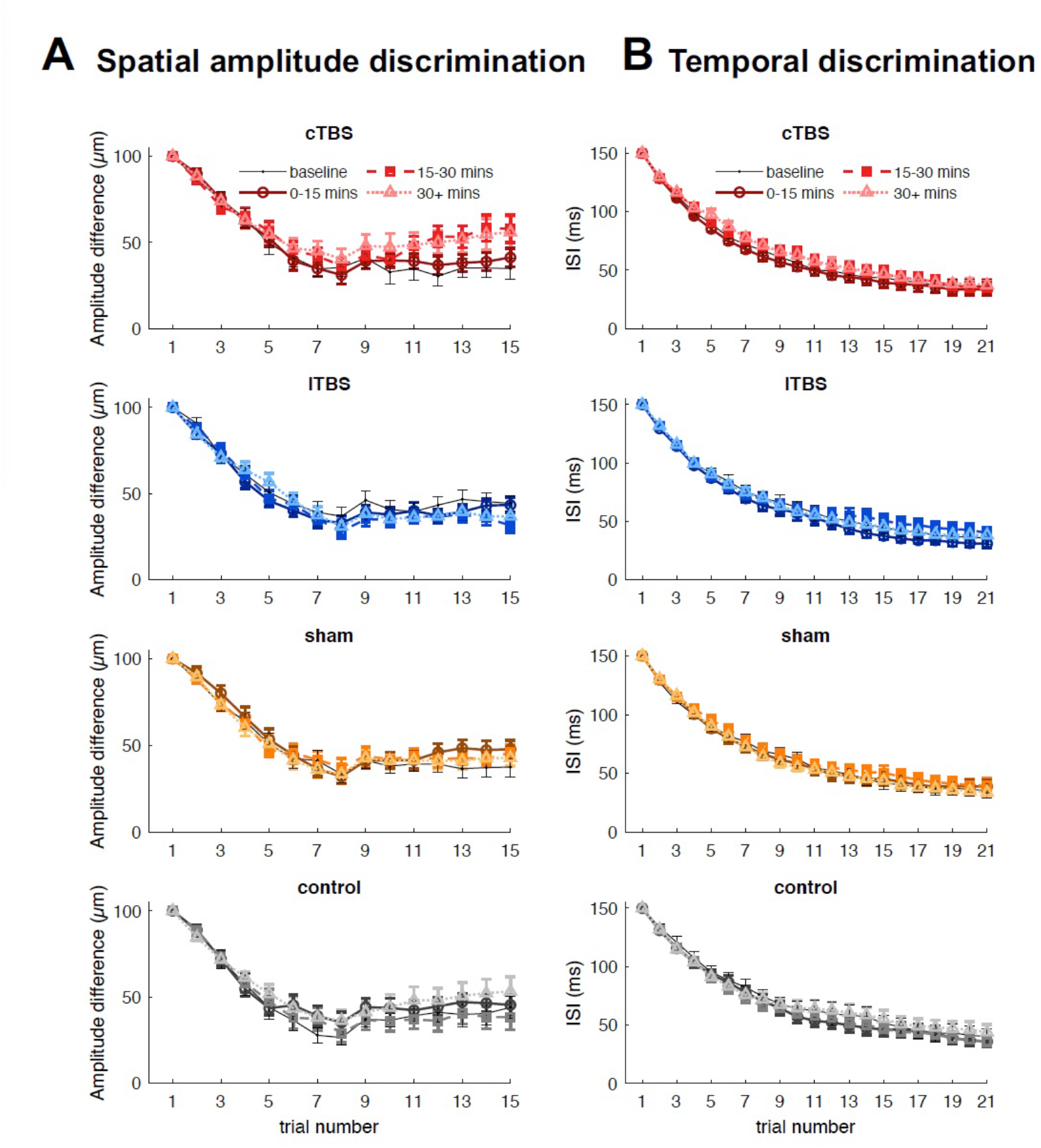
Group-averaged spatial amplitude discrimination performance (A) and temporal discrimination performance (B) for each trial, averaged across blocks within each time window. Lighter colors indicate data from later time windows. Error bars show standard error.

### Temporal discrimination threshold (TDT) task

The mean temporal discrimination thresholds during the baseline phase were 37.3 ± 20.8 ms (cTBS), 37.6 ± 20.3 ms (iTBS), 37 ± 21.4 ms (sham) and 42.4 ± 27.7 ms (no stimulation control), with no significant difference in TDT across sessions (F_(3, 42)_ = 0.34, p = .795, Figure S2). The across-session ICC for baseline TDT was estimated to be 0.77, suggesting a good reliability of the temporal discrimination task (Koo & Li, 2016). In contrast to the effects of cTBS and iTBS on SDT, neither cTBS nor iTBS had effects on TDT (Figure 2B). No significant change was observed during any session (p > .13 in all cases). The baseline-normalized thresholds did not differ significantly between sessions at any post-TBS time window. Group-averaged trial-by-trial temporal discrimination performance for each session is shown in Figure 3B. Performance across all time windows (baseline, 0-15 min, 16-30 min, and 30 min after stimulation) for all sessions reached a plateau around trial 17 of 21.

## Discussion

The current study establishes the capacity of TBS over the tongue representation in S1 to modulate tactile perception of the tongue. Specifically, cTBS was found to impair spatial amplitude discrimination, while iTBS enhanced it. The maximal effects did not occur until 16-30 minutes after the end of TBS. In contrast to the modulatory effects on spatial discrimination, neither cTBS nor iTBS influenced temporal discrimination, suggesting TBS affects the sensitivity, but not timing, of somatosensory afferent signals. TBS thus presents a new way to modulate somatosensation in the orofacial system.

It is noted that the effect of iTBS was smaller than cTBS (+20.08 µm, Cohen’s *d* = 0.7) vs. -8.07 µm, Cohen’s *d* = 0.56). Moreover, although the threshold changes induced by iTBS did differ from the baseline phase, they did not show significant differences when directly compared to control sessions (sham or no stimulation). This is consistent with previous TBS studies on the hand showing that the effect of iTBS over S1 tends to be less robust and stable than that of cTBS (see review, Tang et al., 2023). Spatial amplitude discrimination relies on a combination of excitation within and inhibition between adjacent cortical columns in the S1 (Friedman et al., 2008). In the present study, the application of cTBS may suppress the underlying excitability within S1 cortical columns, leading to a reduction in the lateral inhibition and subsequently affecting the distinction between the two spatially segregated tongue sites. In contrast, iTBS may act to increase the underlying excitability and subsequent lateral inhibition, thereby leading to improved discrimination between the two stimuli. However, it should be noted that the SDT changes observed in our study may be attributed to multiple mechanisms. A previous study found that somatosensory evoked potentials (SEPs) and high frequency oscillations (HFOs) were differently modulated by cTBS and iTBS: iTBS facilitated SEPs without changes in HFOs whereas cTBS modulated HFOs without changes in SEPs (Katayama et al., 2010). This suggests that iTBS and cTBS might affect somatosensory activity and perception through different mechanisms. Such differences could potentially explain the different effect sizes observed following cTBS and iTBS. Future studies are needed to clarify the neurophysiological effects of cTBS and iTBS over S1 and their relationship with behavioral performance.

Interestingly, the maximal modulatory effect did not appear immediately following TBS. Instead, significant increases and reductions of SDT were observed from 16-30 min following cTBS and iTBS respectively. This is in line with Katayama et al. (2010), showing both cTBS and iTBS over S1 caused changes in the excitability of intracortical circuits, as measured by SEPs and HFOs, but these effects were only observed 15-30 minutes after TBS. Our findings, however, are less consistent with a previous behavioral study showing that SDT on the hand was impaired immediately after the application of cTBS and persisted for up to 18 minutes (Rai et al., 2012). It is unclear whether the inconsistency is attributed to participant variability, as responses to TMS can vary largely among individuals, or differences in methodology. For example, in our study, we introduced a 1-minute break after each testing block to prevent tongue fatigue, a design that was not present in the hand study.

In contrast to the modulatory effects of S1-TBS on spatial amplitude discrimination of the tongue, neither cTBS nor iTBS influenced temporal discrimination. Mixed results have been reported from studies investigating the effects of TBS over S1 on temporal acuity of the hand. Lee et al. (2013) observed no changes in temporal acuity after applying cTBS over S1, while other studies found significant impairment following cTBS (Rai et al., 2012) and improvement following iTBS (Conte et al., 2012). Unlike spatial amplitude discrimination, temporal discrimination has been found to engage multiple cortical areas, including the secondary somatosensory cortex (Pons et al., 1992; Romo et al., 2002), anterior cingulate, and supplementary motor areas (Lacruz et al., 1991; Pastor et al., 2004). Moreover, growing evidence suggests that the superior temporal gyrus, prefrontal and parietal cortices might play an important role in tactile temporal perception (Bolognini et al., 2010; Takahashi et al., 2012). Therefore, the lack of change in TDT may be caused by the compensatory involvement of other cortical areas, which could offset the S1 excitability changes induced by TBS. In addition, it should be noted that in the current study, even though participants were given earplugs to minimize the auditory cues resulting from the vibrating probes, some participants might still be able to use these auditory cues during the temporal discrimination task, potentially affecting their performance.

The present study establishes the validity of using neuromodulatory techniques to modulate somatosensation in the orofacial system. Importantly, applying TBS over S1 has the potential to affect all types of orofacial somatosensation, including proprioceptive signals. Due to the inability to block proprioception of the tongue using anesthetics (Putnam & Ringel, 1976; Scott & Ringel, 1971), a lingering question in the field of speech motor control is the impact of a complete loss of proprioception on speech (Parrell et al. 2019). Using TMS to modulate somatosensation in the orofacial system may serve as an improvement over previous attempts to decrease somatosensory acuity through the use of topical anesthetics (Putnam & Ringel, 1976; Scott & Ringel, 1971), which affect only tactile sensitivity. In addition, using TMS to modulate orofacial somatosensation can also benefit clinical populations with abnormal tactile acuity, such as patients with Parkinson’s disease (Chen & Watson, 2017). Therefore, directly modifying somatosensation of the human orofacial system can provide valuable insights into the role of sensory systems in speech production, and has the potential to improve tactile acuity and speech production in both healthy and clinical populations.

## Acknowledgments

Funding: This work was supported by NIH Grant R01 DC019134, a grant awarded through the University of Wisconsin-Madison Fall Research Competition, and a core grant to the Waisman Center from the National Institute of Child Health and Human Development (P50HD105353).

## Author contributions

D-L. T., B.P. and C.A.N designed the experiments. D-L. T. performed research. D-L. T. analysed the data. All authors wrote or edited the manuscript for publication.

## Supplementary materials

**Figure S1:**
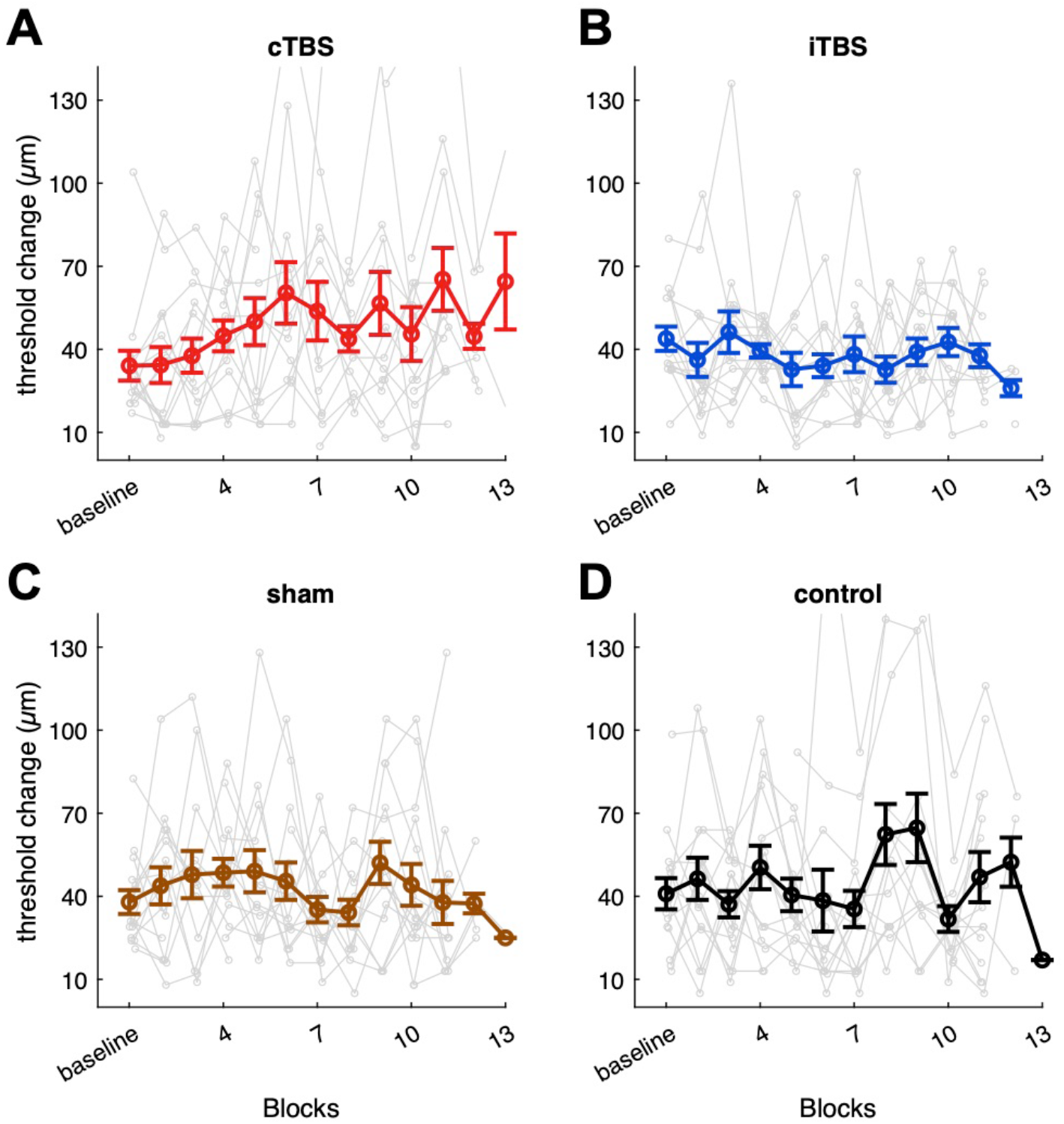
Spatial amplitude discrimination thresholds (SDT) for each block before (baseline) and after stimulations are shown for individuals (small dots, thin lines) and group mean data (large dots, thick lines). Error bars show standard error. A: cTBS stimulation, B: iTBS stimulation, C: sham stimulation, D: control session (no TMS stimulation).

**Figure S2:**
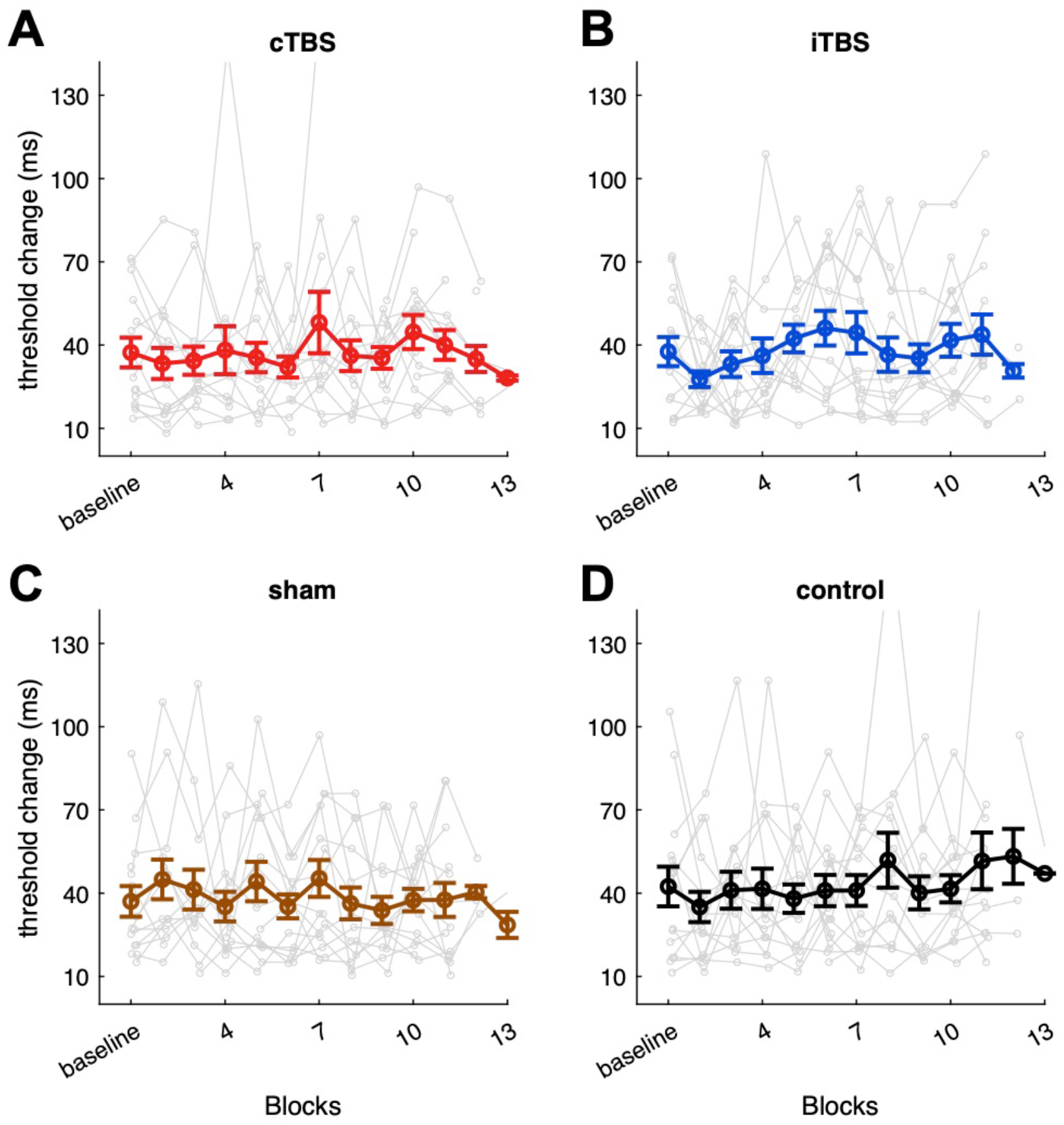
Temporal discrimination thresholds (TDT) for each block before (baseline) and after stimulations are shown for individuals (small dots, thin lines) and group mean data (large dots, thick lines). Error bars show standard error. A: cTBS stimulation, B: iTBS stimulation, C: sham stimulation, D: control session (no TMS stimulation).

